## Supplementary materials for "Oscillatory brain networks in continuous speaking and listening"

| **Speech production** | **Speech production with masked perception** |
|---|---|
| What are your plans for today and the coming days? | Describe a popular artist / author / regisseur. What makes them interesting to you? |
| Which animals do you like? | Describe a traditional christmas. |
| Which hobbies do / did you have? | Where would you like to go on vacation? |
| What does a typical weekend look like for you? | Which places do you like to go in Münster and what is there to see / do? |
| What types of food do you like? | Describe what happens during the olympic vgames. |
| Describe a popular singer / musician / composer. What do you like about them? | Describe the geography and nature of Africa. |
| Which movie-, literature-, comic-character would you like to be and why? | What comes to your mind when you think of poker? |

**Supplementary Table 1:** Full list of questions participants had to answer in the first (speech production) and second recording (speech production with masked perception).

*Speech-brain coupling for listening condition*

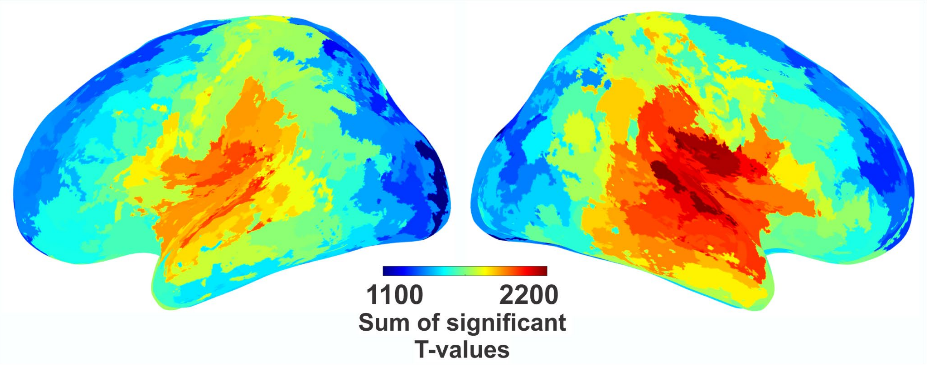


**Supplementary Figure 1: Results of speech brain coupling for the listening condition.** Cortical map represents groups statistics of multivariate MI between speech envelope and brain activity in each parcel compared to 95th percentile of surrogate data. Colour code represents the sum of all significant T-values across delay and frequency (FDR-corrected across delays (-1-1 s), frequencies (1 to 10 Hz) and parcels).

*Comparison of our speech production network with fMRI meta-analysis*

To assess the localisation accuracy of our MEG speech production network based on speech-brain coupling we compared our results to the automatic meta-analysis provided by neurosynth.org. Using the term ‘speech production’ resulted in a meta-analysis of 107 fMRI studies investigating speech production. We downloaded the statistical meta-analysis map that corresponded to a uniformity test. The map of z-scores results ‘from a one-way Anova testing whether the proportion of studies that report activation at a given voxel differs from the rate that would be expected if activations were uniformly distributed throughout gray matter’ (quote from neurosynth.org). The map therefore corresponds to a classical fMRI statistical analysis and measures consistency of activations across studies. This volumetric statistical map was then transformed to the HCP atlas in the following way: First, we interpolated the statistical map to the surface representation of the HCP atlas using ft_sourceinterpolate.m in fieldtrip. Next, we averaged all statistical values within the same anatomical parcel. This resulted in one value per parcel that could be directly compared to the statistical maps from our study (which are also based on one value per parcel). Both statistical maps represent a measure of consistency (across studies for neurosynth and across participants for our study) and can be expected to be similar. We therefore hypothesised that parcels with high statistical values in the fMRI meta-analysis (neurosynth) also show high statistical values in our study (and correspondingly for parcels with low values). We therefore correlated the statistical values of both statistical maps across parcels. The correlation was highly significant ($r=0.42$, $p<<0.0001$). To further test significance we performed a permutation test and computed the correlation 1000 times on randomly permuted values which resulted in a 99th percentile of this null distribution of $r_0=0.16$. Both results demonstrate a highly significant relationship between both statistical maps and indicate that our results are valid.

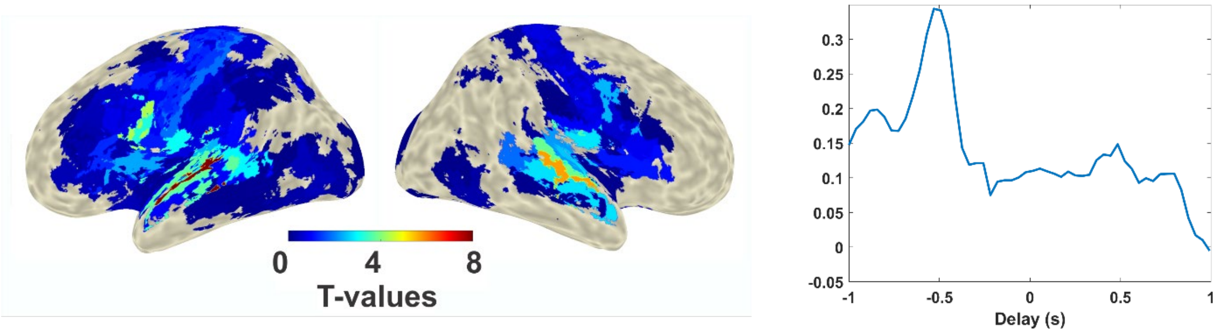


**Supplementary Figure 2:** Upper panel**:** Statistical map of fMRI meta-analysis from neurosynth.org using the search term ‘speech production’**.** Results were spatially interpolated to the HCP atlas used in our study. Color codes correspond to z-values from a uniformity test as documented on neurosynth.org. Lower Panel: The relative difference of correlations (r1-r2)/(r1+r2) across delays. r1 is the correlation between our speech production network and the neurosynth speech production network and r2 is the correlation between our speech production network and the neurosynth speech perception network.

Still, one could argue that correlations between our statistical map and the speech production meta-analysis from neurosynth.org might be high and significant for any network and therefore not specific for the speech production network. Therefore, we repeated the analysis for three other networks: motor network, resting-state network and speech perception network (the names used here also represent the neurosynth search terms). We like to note that including the speech perception network represents a formidable challenge since it shows a large overlap with the speech production network. As before, we correlated these three additional maps with our speech production map (Figure 2) and observed the following correlations (speech perception $r=0.33$, $p<<0.001$; motor network $r=0.18$, $p=0.007$, resting state $r=-0.07$, $p>0.05$, all $df=228$). Therefore, our speech production network most closely resembles the speech production network from neurosynth. To test how robustly the correlation with the neurosynth speech production network is higher than the neurosynth speech perception network we performed 10000 bootstrap correlations (with random selection of 230 parcels with replacement in both statistical maps) and counted how often the correlation with the speech production network was higher than the correlation with the speech perception network. This was the case in 99.7% of the bootstrap iteration indicating that our network robustly resembles the neurosynth speech production network better than the neurosynth speech perception network.

Finally, in another stringent test of the validity of our results we tested the hypothesis that the higher correlation of our speech production network to the neurosynth speech production network compared to the neurosynth speech perception network is delay dependent. Specifically, the correlation should be highest for negative delays (where brain activity precedes the speech envelope. The lower panel of Suppl. Figure 2 shows the relative difference of correlations (r1-r2)/(r1+r2) across delays. r1 is the correlation between our speech production network and the neurosynth speech production network and r2 is the correlation between our speech production network and the neurosynth speech perception network. A clear positive peak is evident at negative lags indicates that the best match between speech production networks in neurosynth and in our analysis occur for brain activity preceding the speech envelope.

*Lateralisation (Right vs left) for listening*

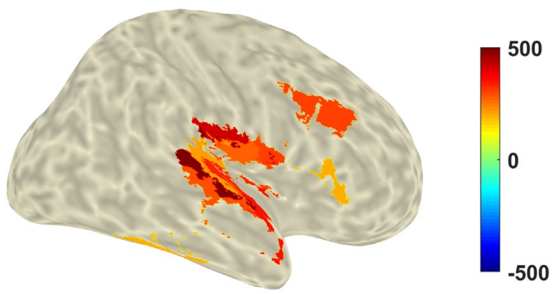


**Supplementary Figure 3: Speech-brain coupling lateralisation for listening condition.**

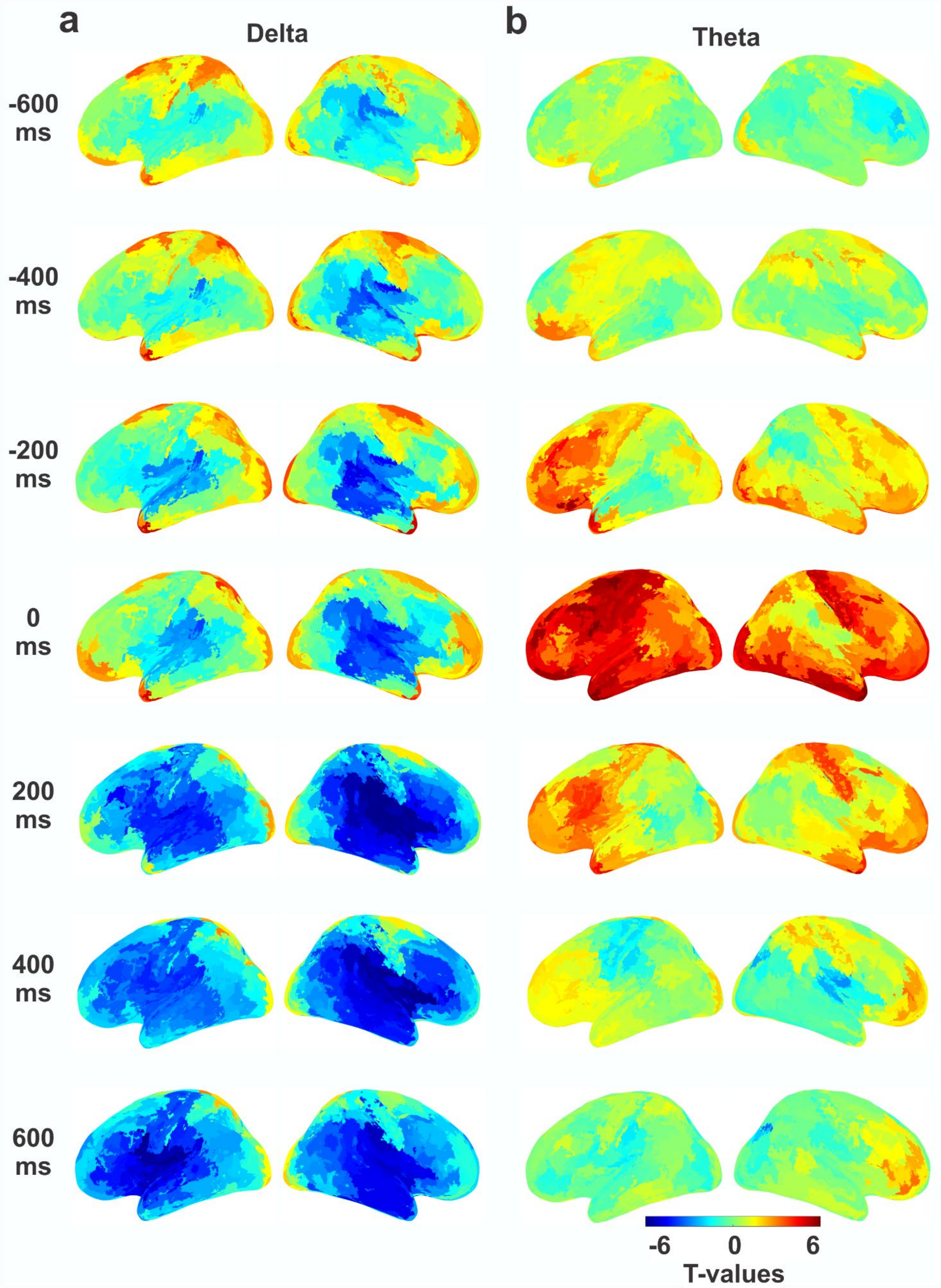


**Supplementary Figure 4:** Statistical comparison of speech-brain coupling in speaking versus listening at delta (2 Hz) and theta (5 Hz) for different delays ([-600 600] ms). Colour codes t-values.

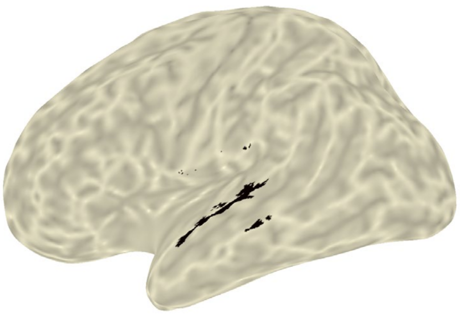

**Supplementary Figure 5: Left STG area.** The highlighted black parcel depicts the L_A5 parcel of the HCP atlas representing the LSTG area.

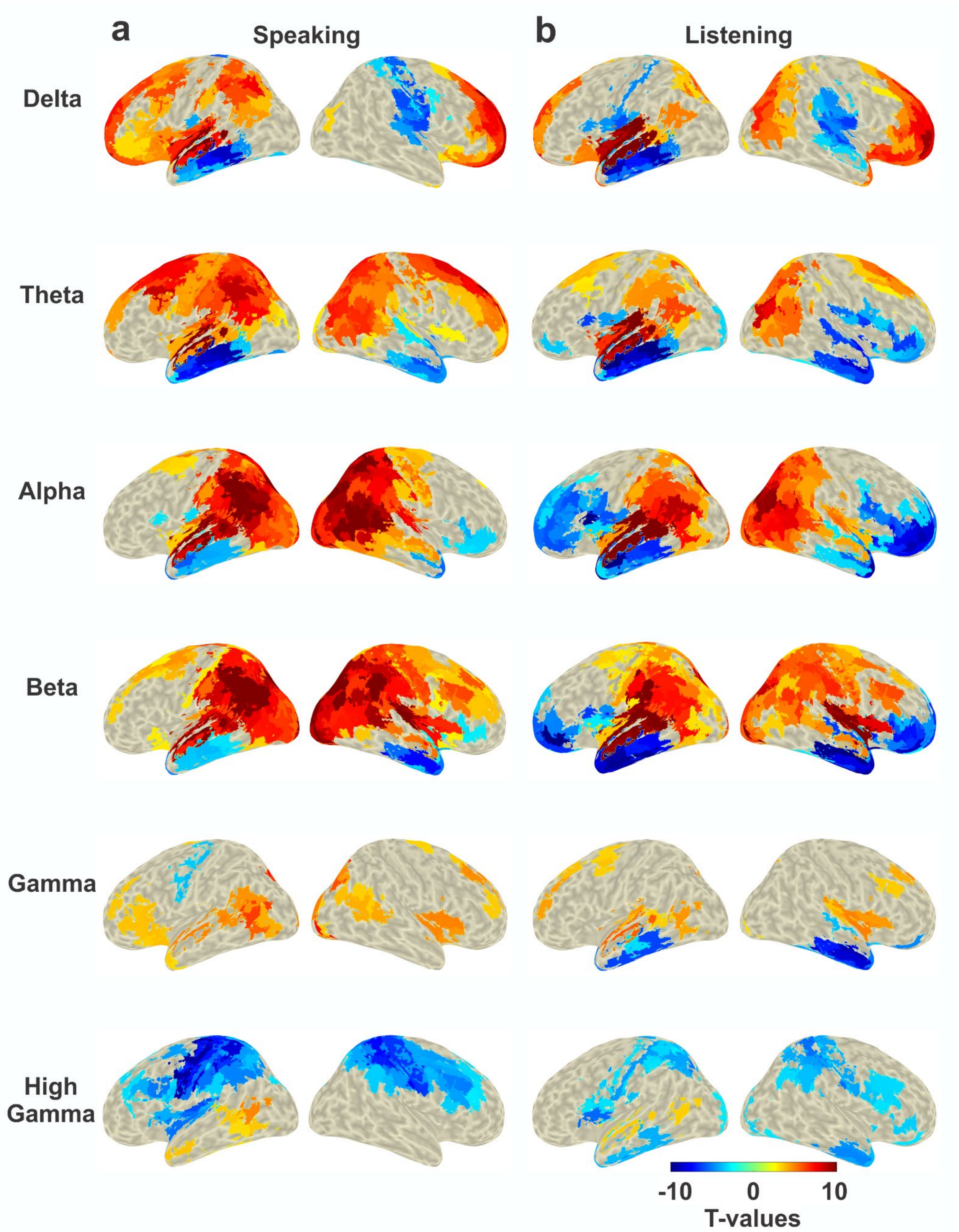


**Supplementary Figure 6:** Significant connectivity between STG and other cortical parcels during speech production (a) and listening (b) in different frequency bands. A cluster-based permutation test was used to detect significant connectivity patterns ($p<0.05$). Colour codes t values. Blue colour represents the flow of information from STG to other cortical parcels and the red colour represents the opposite direction.

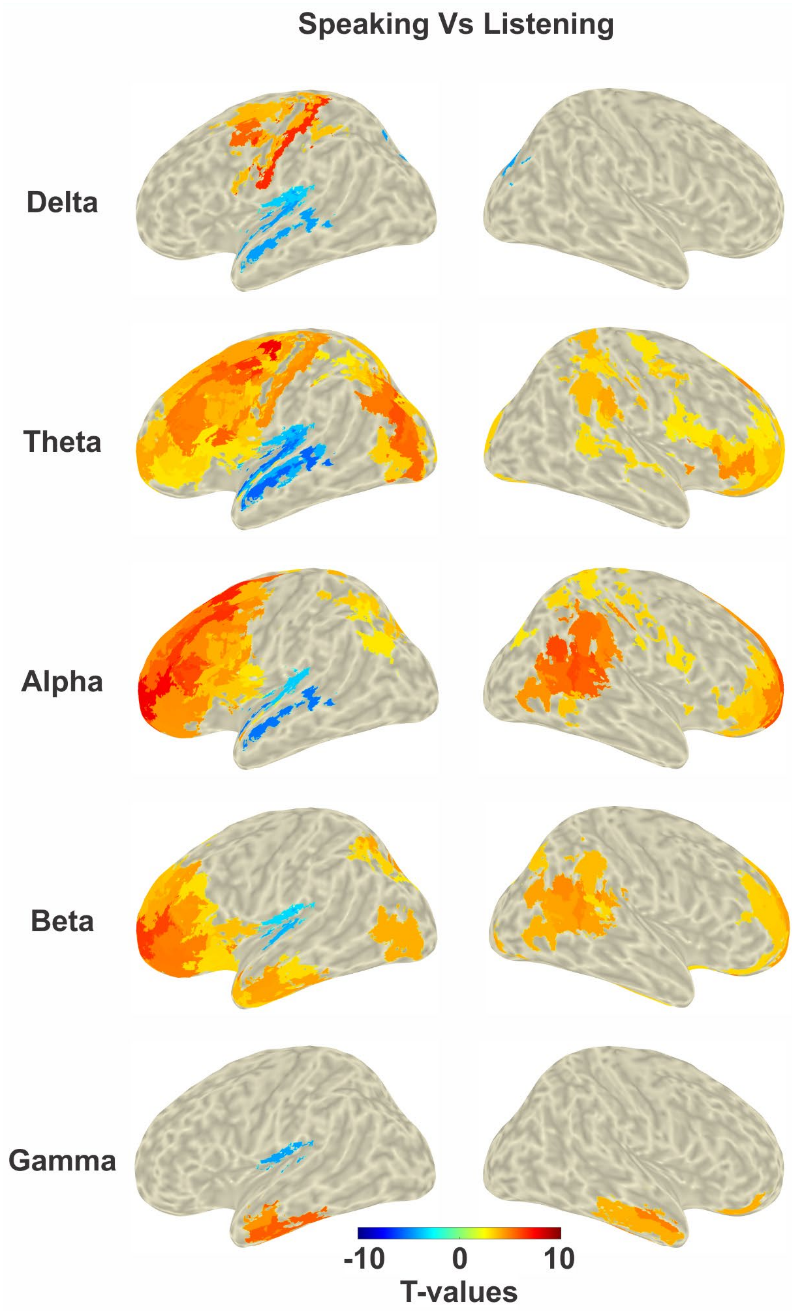


**Supplementary Figure 7:** Comparison of connectivity patterns between speaking and listening conditions in different frequency bands. A cluster-based permutation test was used to detect significant connectivity differences from all the cortical parcels to STG between speaking and listening conditions ($p<0.05$). Colour codes t values. Please note that since we compared asymmetry indices (DAI) between two conditions, interpreting the directionality from this cortical plot is challenging. For a better understanding of these statistical maps, please refer to the spectrally-resolved DAI between STG and four ROIs in figure 7d as well as supplementary figure 6.

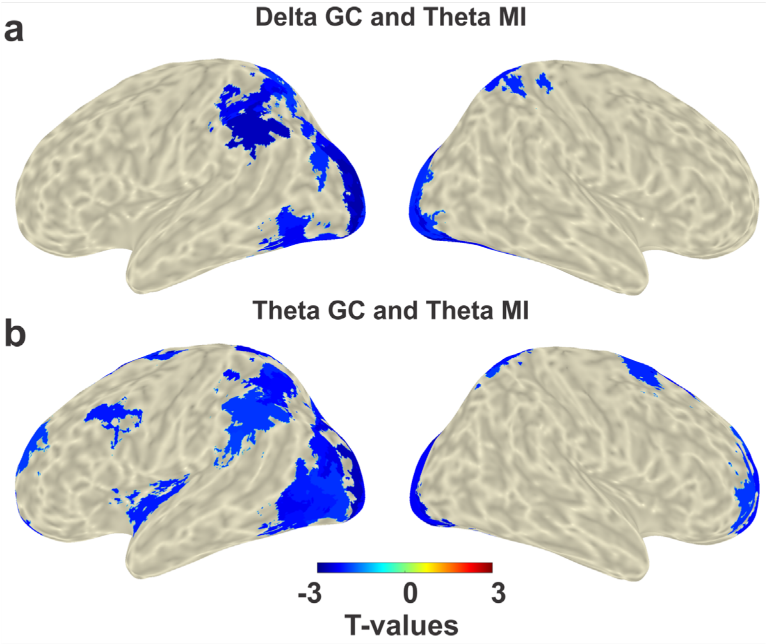


**Supplementary Figure 8: Correlation analysis between top-down GC and MI.** Speech-STG coupling in theta range (positively lagged: 130ms) is negatively correlated with the top-down delta (a) as well as theta (b) connectivity from mainly left occipital and parietal areas.

*Statistical contrast of frequency-specific power between speaking and listening*

We also calculated the statistical contrast of frequency-specific power (not coupling to the speech envelope) between speaking and listening. The analysis reveals significant positive and negative effects. However, positive effects (higher power in speaking compared to listening) could be (partly) caused by residual speech artefacts (such as muscle artefacts or movement). Therefore, in Supplementary Figure 1 we report only effects where power in any frequency band is significantly lower in the speaking condition compared to listening. As expected, we see the strongest power decrease (during speaking compared to listening) over bilateral motor areas extending to premotor and parietal areas with a dominance in the left hemisphere. The power suppression in motor and premotor areas includes medial supplementary motor areas (SMA) and extends all the way down to lateral motor areas corresponding to speech-related motor representations of larynx, tongue, and mouth. Supplementary Figure 1b shows that these suppression effects arise from power differences at frequencies between about 10 Hz and 30 Hz, largely corresponding to the beta frequency band that has been consistently related to motor functions [89,90].

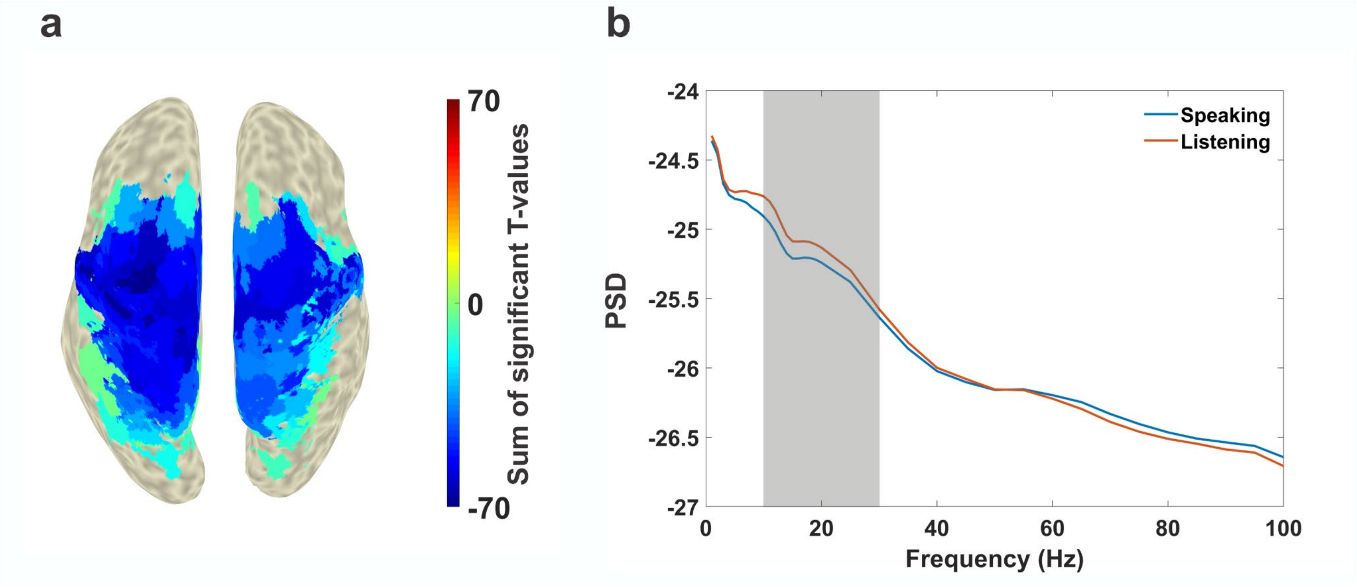


**Supplementary Figure 9. Frequency-specific power comparison between speaking and listening condition. a.** The statistical contrast of frequency-specific power between speaking and listening. Results of dependent-samples t-test were FDR corrected across parcels and frequencies (1-100 Hz). Colour codes sum of t-values across all significant frequencies. Note that we use a one-tailed test (speaking < listening) to ensure that results are not contaminated by residual speech artefacts. **b.** Power spectral density averaged across all significant parcels for speaking (blue) and listening (red) conditions. The grey panel depicts the frequency range 10-30 Hz.
